# Resolving Complex Structural Genomic Rearrangements using a Randomized Approach

**DOI:** 10.1101/028217

**Authors:** Xuefang Zhao, Sarah B. Emery, Bridget Myers, Jeffrey M. Kidd, Ryan E. Mills

## Abstract

Complex chromosomal rearrangements consist of structural genomic alterations involving multiple instances of deletions, duplications, inversions, or translocations that co-occur either on the same chromosome or represent different overlapping events on homologous chromosomes. We present SVelter, an algorithm that first identifies regions of the genome suspected to harbor a complex event and then iteratively rearranges the local genome structure, in a randomized fashion, with each structure scored against characteristics of the observed sequencing data. We show that SVelter is able to accurately reconstruct these regions when compared to well-characterized genomes that have been deep sequenced with both short and long read technologies.

## BACKGROUND

Structural variation (SV), defined as chromosomal rearrangements resulting from the removal, insertion or rearrangement of relatively long regions of the genome, are natural sources of genetic variation[1-3] that have also been implicated in numerous human diseases including neurological disorders[4-6]. There have been extensive studies to discover these genomic aberrations from the whole genomes of humans and other species, and numerous algorithms have been developed to accurately identify their prevalence[7-11]. These approaches have primarily focused on simple copy number variant (deletions, duplications) or copy neutral (inversions) rearrangements defined by at most two chromosomal breakpoints and work by identifying and clustering various signals of discordant alignments from paired-end next generation sequencing data[12] such that deletions are inferred from read pair clusters with abnormally long apparent insert length or reduced depth of sequence coverage while simple inversions can be detected with clusters of read pairs with aberrant orientation. Recent algorithms have begun to integrate signals across multiple features to increase sensitivity[9, 11, 13] and these have been successful in precisely identifying various types of SVs. Knowledge of the underlying structure of the rearrangement is still required, however, in order to properly model these aberrant signals to the correct type of structural variant. For example, clusters of read pairs with insert sizes larger than expected are typically representative of deleted sequence since this observation is consistent with how the reads would map in the presence of such an event.

Recently, more complex SVs (CSVs) consisting of multi-breakpoint or overlapping genomic rearrangements have been revealed to be more pathologically significant than previously thought[4, 5, 14] and have been either neglected or misinterpreted by current techniques due to the complexity of the signals shown by the sequencing data. This is primarily due to the limitations of presupposing the types of SVs being considered, as often times the signals from one event are clustered independently from those of another and can lead to contradictory or sometimes even opposing predictions to what is actually present. Under such circumstances, traditional prediction models lose their ability to discriminate between signals, and therefore new computational strategies are required to overcome these challenges.

Here, we present a novel approach, SVelter, to accurately resolve complex structural genomic rearrangements. Unlike previous ‘bottom up’ strategies that search for deviant signals to infer structural changes, our ‘top down’ approach works by virtually rearranging segments of the genomes in a randomized fashion and attempting to minimize such aberrations relative to the observed characteristics of the sequence data. In this manner, SVelter is able to interrogate many different types of rearrangements, including multi-deletion? and duplication-inversion-deletion events as well as distinct overlapping variants on homologous chromosomes. The framework is provided as a publicly available software package that is available online (https://github.com/millslab/svelter).

## RESULTS

### Overview of SVelter

Our approach predicts the underlying structure of a genomic region through a two-step process. SVelter first identifies and clusters breakpoints (BP) defined by aberrant groups of reads that are linked across potentially related structural events. It then searches through candidate rearrangements using a randomized iterative process by which rearranged structures are randomly proposed and scored by statistical models of expected sequence characteristics (Figure 1; Materials and Methods).

**Figure 1.**
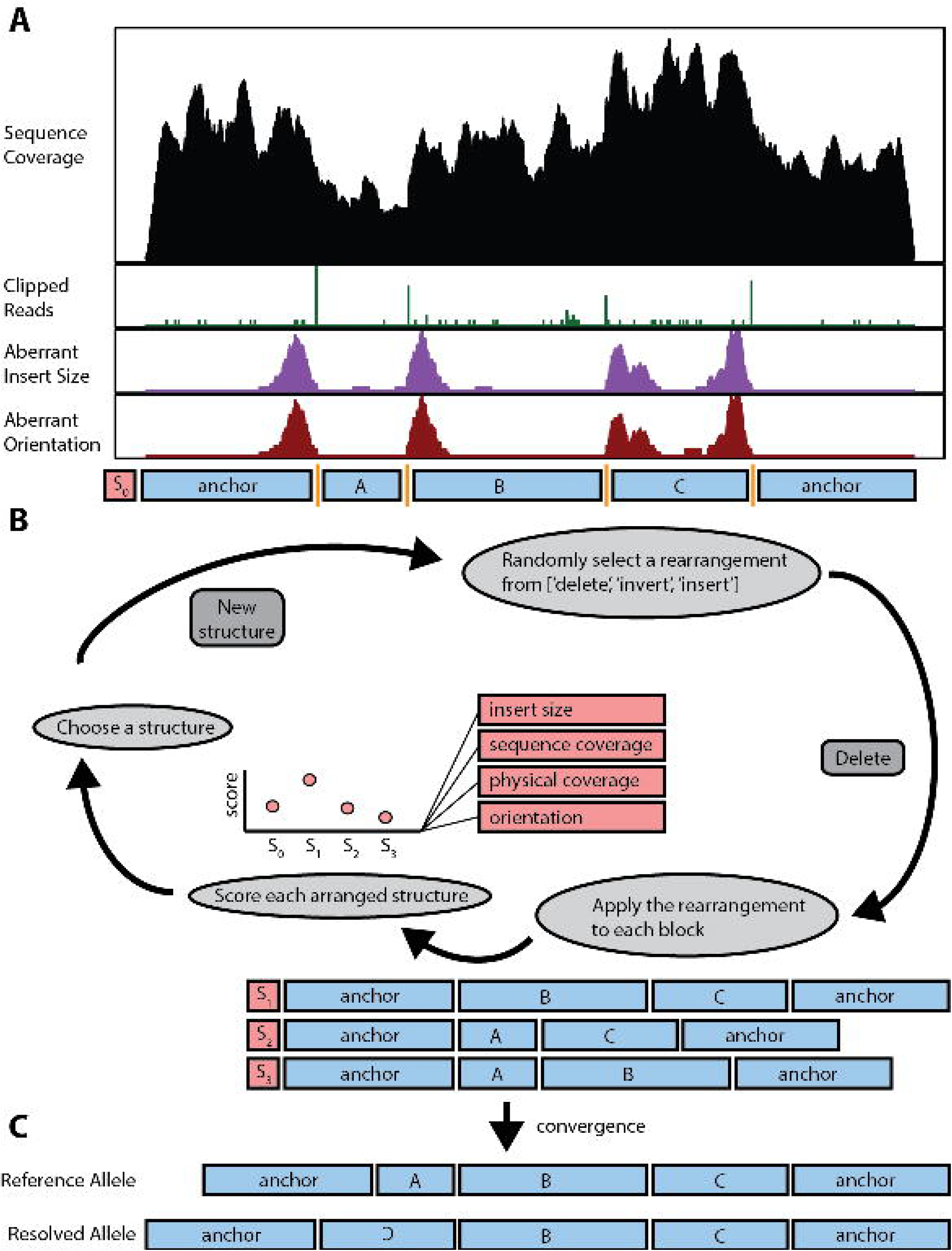
Overview of computational strategy for identifying structural variation in whole genome sequences. **(A)** SVelter first scans the genome and identifies clusters of aberrant read characteristics. These are used to create a putative set of breakpoint positions. **(B)** The segments between breakpoints are then iteratively rearranged and scored against null models of sequence expectations. **(C)** The final converged structure is reported as the predicted structural rearrangement for the region.

SVelter begins by fitting statistical models for insert size (IS) and read depth (RD) based on paired-end sequences sampled from copy neutral genomic regions[15]. Both are modeled as normal distributions for efficiency purposes which is recommended for relatively clean data sequenced at higher depth; however, more accurate but slower models (i.e. binomial for IL and negative binomial for RD) are also available as options for data of lower quality. SVelter then searches for and integrates potential SV signals from read pairs with aberrant insert size, orientation, and/or alignment (soft-clipping). Pairs of BPs are assigned simultaneously, and BP pairs that intersect with each other are further connected to form BP clusters. For each cluster containing *n* BPs, the *n-1* genomic segments defined by adjacent BPs are then rearranged in a randomized iterative process whereby a simple SV (deletion, insertion, inversion) is randomly proposed and applied to all possible segments to assess the viability of each putative change. The initial structure and each subsequent rearranged structure are then scored by examining the impact of each change on various features of the sequence reads in the region, including insert size distribution, sequence coverage, physical coverage, and the relative orientation of the reads. A new structure is then chosen for the next iteration using a probability distribution generated from the structure scores. This continues until the algorithm converges on a final, stable set of rearrangements or a maximum number of iterations is reached.

An important feature of SVelter is that it simultaneously constructs and iterates over two structures, consistent with the zygosity of the human genome. This allows for the proper linking of breakpoint segments on the correct haplotypes, which is crucial for the proper resolution of overlapping structural changes that can often confuse or mislead other approaches. Individual breaks in the genome can then be properly linked and segregated, aiding in downstream genotyping across multiple individual sequences.

The randomized aspect of this approach leads to additional computation cost relative to other SV detection algorithms. We have addressed this by implementing a number of optimizations to increase the overall efficiency of SVelter. First, we limit the number of clustered BPs during the initial breakpoint-linking step in order to manage the number of random combinations at the next step. For regions with potentially higher numbers of linked breakpoints, we form subgroups based on physical distance between adjacent BPs that are later combined. Second, we set an upper and lower bound on the potential copy number (CN) of each segment between BPs using local read depth information and only allow structures containing *CN-1* to *CN+1* blocks for downstream analysis. Lastly, we have restricted the total number of iterations such that after converging on a stable rearrangement for 100 continuous iterations, we set this structure aside and restart the random iterations for another 100 iterations, at which point the highest scoring structure overall is chosen. This results in a total processing time for SVelter on a re-sequenced human genome with 50X coverage of under 20 hours when run in parallel on a high-performance computing cluster.

### Performance Evaluation

We compared SVelter to three popular SV detection algorithms: Delly[11], Lumpy[9], and Pindel[8]. Both Delly and Lumpy have integrated insert size and split read information into their SV detection strategy, while Pindel implements a pattern grown approach to utilize split read alignments. While there are numerous other algorithms that have been developed for detecting SVs, we focused on these as they have previous published comparisons that can be transitively applied to our results.

Multiple experiments were conducted in order to evaluate our approach. We first simulated genomes of various sequence coverage containing both simple and complex SVs as homozygous and heterozygous events. We next applied these algorithms to the genome of a haploid hydatidiform mole (CHM1)[16] and also a well-characterized diploid genome (NA12878)[17, 18], both of which had reported high-confident calls as well as long-read Pacific Biosciences (PacBio) sequences available for orthogonal assessment. All algorithms were run either with the recommended settings as provided by the authors (where available) or default settings. Detailed commands for running each algorithm can be found in supplemental materials.

#### Simulated data

We simulated heterozygous and homozygous non-overlapping simple SVs (deletions, inversions, tandem duplications, dispersed duplications and translocations) of varied sizes into synthetic genomes sequenced at different depths of coverage (10-50X). We then calculated the sensitivity and positive predictive value (PPV) of each algorithm (Figure 2A,B, Supplemental Figures 1-2). Overall, SVelter achieves a higher sensitivity and PPV for simple deletions compared to all other algorithms. Comparisons with duplications were more difficult; while all compared approaches can report tandem duplications, for dispersed duplications only SVelter reports both the duplicated sequence and its distal insertion point. We therefore took a conservative approach such that for calculating sensitivity we compared the full set of duplications predicted from each approach to the simulated set of tandem and dispersed events, but limited the false positive analysis to only tandem duplications for the other algorithms. It should be noted that this method of comparison would bias against SVelter to some extent, however under these circumstances SVelter still showed very good sensitivity and positive predictive value in calling dispersed duplications, with slightly worse performance for tandem duplications. For inversions, SVelter showed a comparable accuracy to other the algorithms.

**Figure 2.**
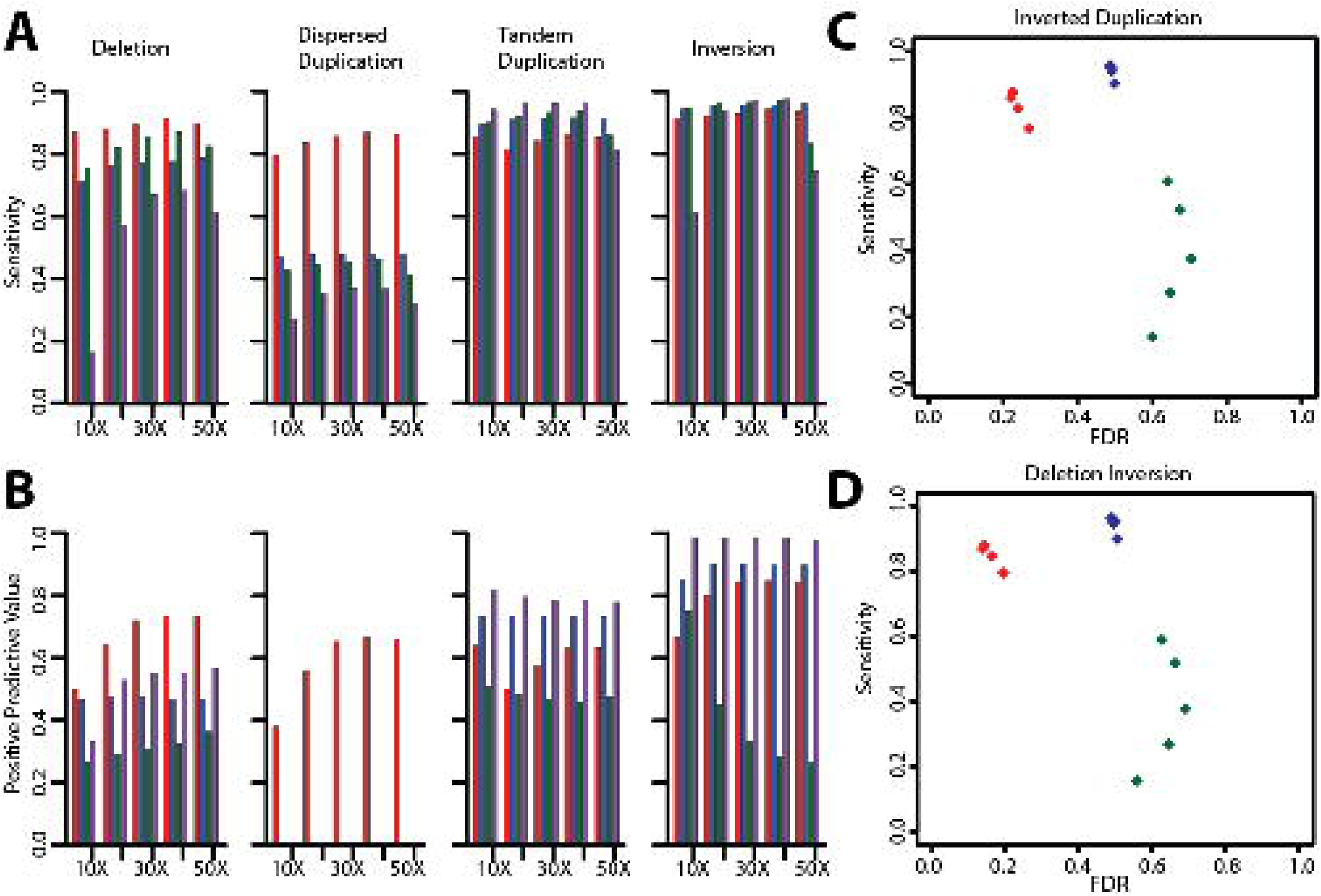
Assessment of accuracy using simulated data sets. **(A)** Sensitivity and **(B)** positive predictive values for SVelter (red), Delly (blue), Lumpy (Green), and Pindel (purple) across different simple SV types and sequence coverage on combined simulated homozygous and heterozygous events. For dispersed duplications, only SVelter was considered for positive predictive values and all predictions by other algorithms that did not overlap simulated results were considered only for the tandem duplication category. **(C)** Sensitivity and false discovery rate (FDR) for simulated complex inverted duplications. **(D)** Sensitivity and FDR for simulated complex inversion deletion events.

We also simulated specific types of complex rearrangements based on structures recently reported[19] as well as our own observations (Supplemental Table 1). Performance comparisons with complex structures are less straightforward than with simple SVs as most algorithms are only designed to identify simple events, and therefore may predict portions of CSVs as independent events. We address this issue by considering the identification and predicted copy number of individual junctions as reported in the entire prediction set of each algorithm (deletions, duplications, inversions) and compared against each simulated complex event collectively, treating predicted non-simulated junctions in the region as false positives (Methods and Materials). SVelter performs consistently better in terms of sensitivity and PPV across almost all types of complex events, including inverted duplications and inversion deletion events (Figure 2C,D).

**Table 1.** Predicted SV Types in CHM1 and NA12878 by SVelter.

| SV Type | CHM1 | NA12878 |
| --- | --- | --- |
| Simple DEL | 1890 (0.86) | 2988 (0.84) |
| Simple DUP | 1872 (0.28) | 1232 (0.41) |
| Tandem | 1725(0.27) | 1184 (0.41) |
| Dispersed | 147 (0.37) | 48 (0.40) |
| Simple INV | 14 (0.50) | 106 (0.76) |
| Simple TRA | 6 (0.67) | 3 (0.67) |
| DEL+DUP | 6 (0.50) | 39 (0.49) |
| DEL+INV | 5 (0.40) | 16 (0.44) |
| DEL+TRA | 3 (0.67) | 3 (0.67) |
| DUP+INV | 188 (0.64) | 141 (0.44) |
| DEL+DUP+INV | 8 (0.50) | 34 (0.32) |
| Other | 27 (0.37) | 369 (0.70) |

Numbers in parenthesis indicate percentage of calls with direct PacBio validation support

#### Real data

To estimate how SVelter performs on real data, we have applied each algorithm to two publicly available datasets: a haploid hydatidiform mole (CHM1)[16] and a well-characterized diploid genome analyzed by the Genome in a Bottle Consortium (NA12878)[17, 18]. Both have been deep sequenced by Illumina short-insert and PacBio long-read sequencing, and provide an excellent foundation for comparing baseline accuracies among approaches. We initially compared deletion calls of each algorithm to the reported set of variants to determine their relative accuracy, however the false discovery rate of each algorithm was abnormally high with respect to previously reported values (Supplemental Table 2), suggesting that the reported deletion set may be overly conservative. We therefore examined the PacBio data directly for each predicted variant using a custom validation approach that utilizes a recurrence strategy to compare each read to both the reference allele as well as a rearranged reference consistent with the predicted structure (Figure 3A,B, Methods and Materials). We evaluated this approach using sets of reported deletions in these samples as well as matched random sets located within copy neutral regions and found it to have very high true positive and true negative rates (Figure 3C). We also conducted PCR experiments on the predicted breakpoints of three predicted complex rearrangements that were validated with this approach to show convincing evidence for two, with inconclusive results for the third due to high degrees of repetitiveness in the region (Supplemental Figures 3-6). We then reassessed the earlier deletion predictions made by each algorithm in CHM1 and NA12878 by combining the previously reported deletions in each sample with those having PacBio validation support from our analysis. As expected, we observed a marked increase in accuracy for each algorithm (Figure 3D).

**Figure 3.**
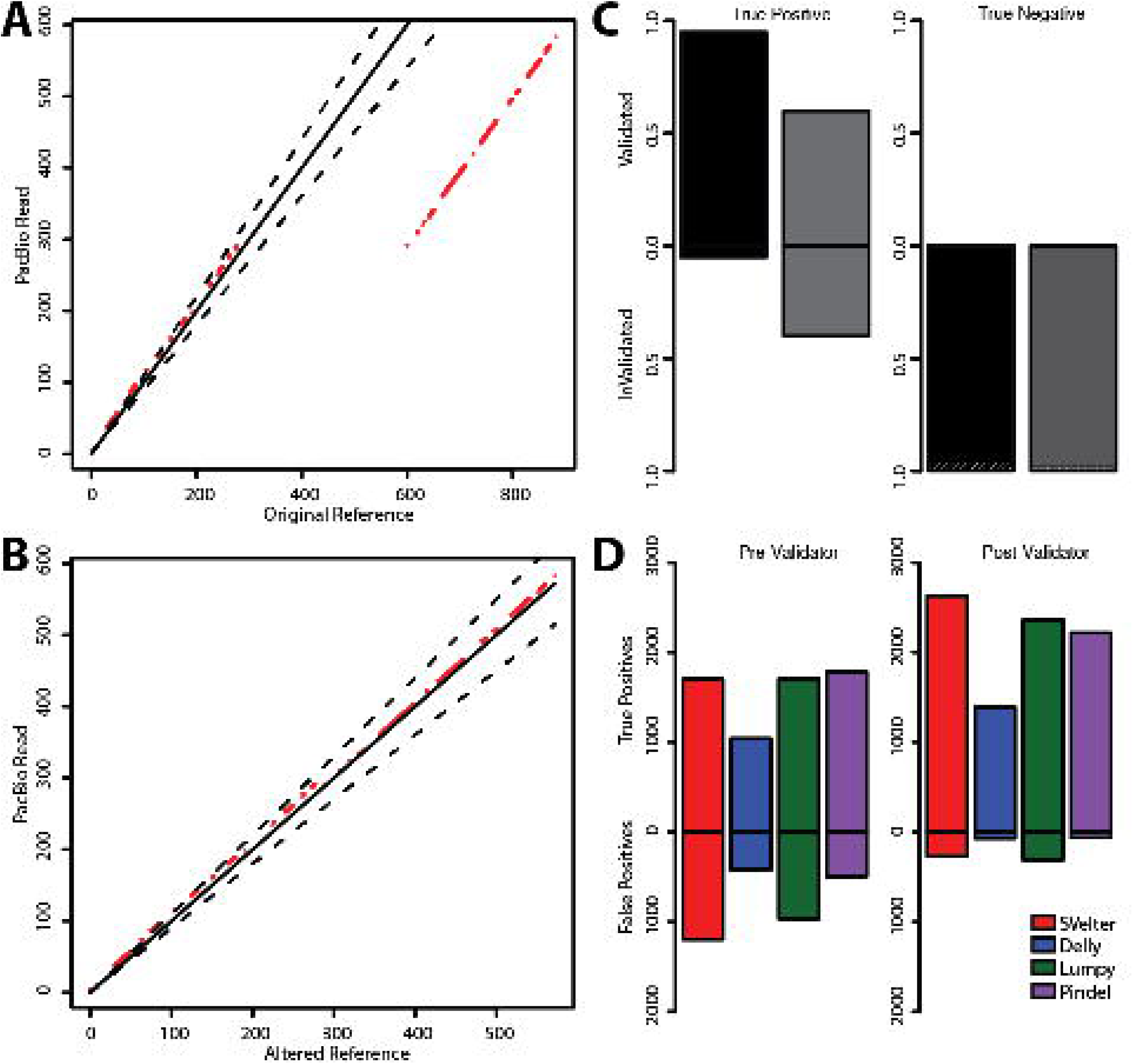
Overview and application of PacBio validation approach to human data. **(A)** Dot plot of example region containing a simple deletion using a single PacBio read against the reference genome. Red dots indicate matches between sequences and dashed black lines delineate 10% deviance from the diagonal. **(B)** Dot plot of same region using an altered reference incorporating the deletion event. **(C)** Fraction of true positive calls using validation approach on published deletions in NA12878 (black) and CHM1 (grey) and CN2 regions as negative controls. Dashed black lines indicate regions that could not be assessed due to lack of PacBio reads to interrogate. **(D)** Performance comparison of deletions predicted by each algorithm using published deletions in NA12878 (left panel) and published deletions combined with PacBio validated calls (right panel).

We next compared the performance of each algorithm on identifying and resolving CSVs. Given that there are very few reference sets available of known complex rearrangements, we first created a set of non-overlapping candidate CSVs as identified by SVelter in CHM1 and NA12878. We then collected all predictions from each algorithm that overlap that region and scored them using the validation approach above. As many complex rearrangements may be described as a combination of simple SVs, we utilized a ranking approach to compare the individual predictions by assigning 0 to the lowest scores and 0.75 to the highest scores (see Methods and Materials). We observed a significant enrichment of SVelter predictions with high validation scores, indicative of its efficacy in correctly resolving CSVs (Figure 4A). An example is shown in Figure 4B, which depicts a summary of sequence read alignments for a region on chromosome 1 in CHM1 containing multiple deletions as well as a local translocation. Using standard read clustering algorithms, the signals present might suggest the presence of a tandem duplication overlapping with large deletions. However, this is not consistent with the haploid nature of CHM1, and comparisons with long PacBio sequence reads that overlap the region show the true structure (Figure 4C), which when aligned to a rearranged reference using SVelter predictions shows a full length alignment (Figure 4D). A comparison with other algorithms indicates that their predictions are indeed consistent with analyzing each aberrant read cluster independently of each other and result in a combination of tandem duplications, deletions, and inversions (Figure 4E).

**Figure 4.**
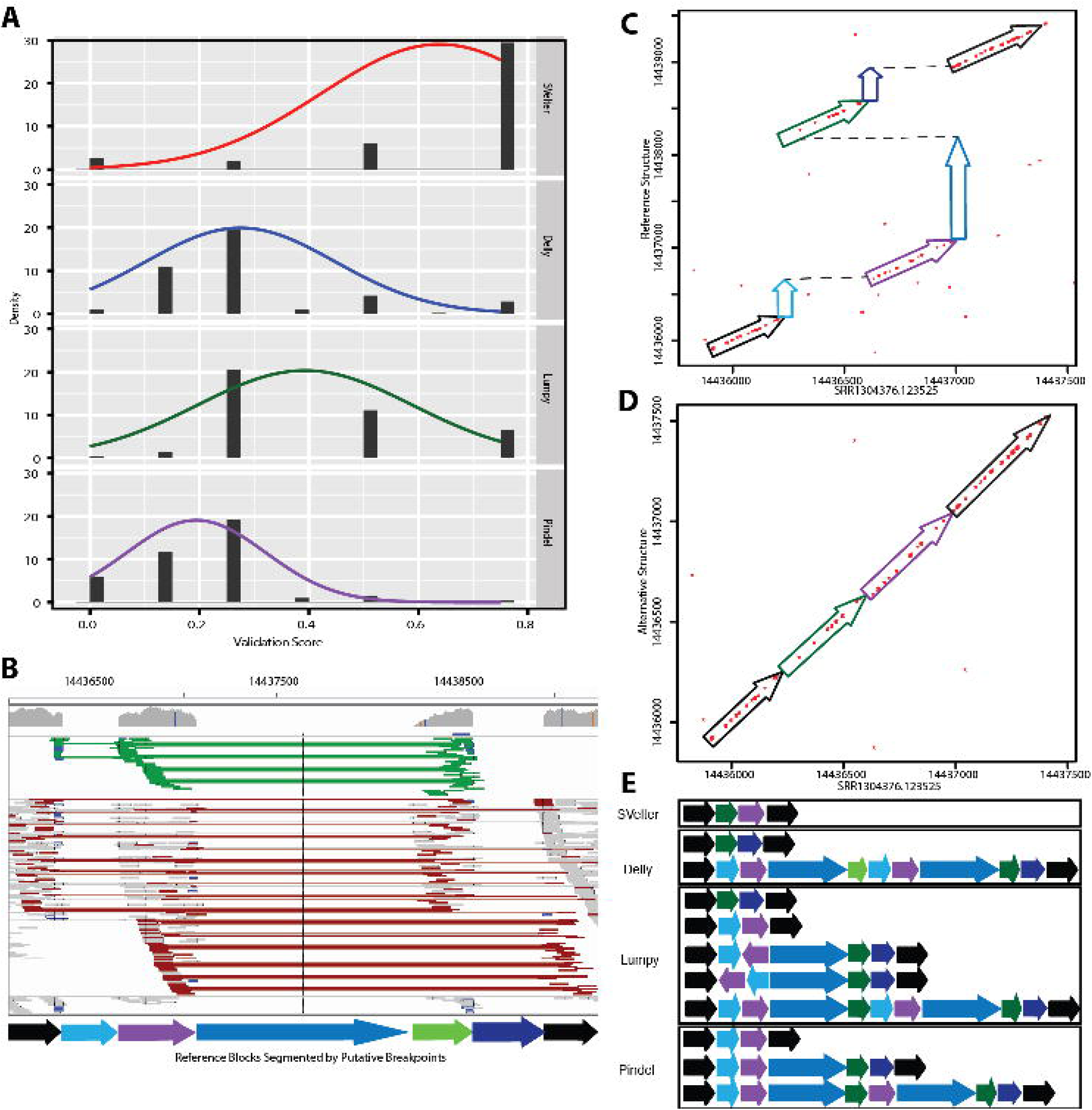
Evaluation of complex structural variation predictions. **(A)** Validation scores of complex structural variation predicted in NA12878 from all algorithms ranked and normalized from 0 to 1 for comparison. For approaches with multiple predicted SVs in a region, average scores from each prediction were averaged. **(B)** IGV screenshot of example complex region in CHM1 (chr1:14435000–1444000) containing multiple deletions (blue shaded arrows) and a translocated region (green arrow), with surrounding anchor regions in black. Light green lines in IGV indicate read pairs with reverse-forward orientation, while red lines indicate read pairs with aberrant insert size length. **(C)** Dot plot of region between an individual PacBio read (SRR1304376.123525) against the reference sequence. Colored arrows correspond to segments indicated in (B). **(D)** Dot plot of altered reference sequence implementing predicted rearrangements by SVelter. **(E)** Schematic of predictions by each SV algorithm with respect to segments indicated in (B). For approaches with multiple predictions overlapping the region, each predicted SV is show independently.

#### Computational Runtime

We compared the overall executable runtime of the different software packages using a single chromosome from NA12878. For each algorithm, we initialized the analysis using a previously aligned sequence in BAM format and used the respective procedures necessary for each approach to result in a variant call file (see Methods and Materials). Delly was observed to complete the fastest, followed by Lumpy. Pindel and SVelter were both considerably slower and were comparable in their runtime (Supplemental Table 3). It should be noted that some algorithms (e.g. Lumpy) can perform faster with optimized alignment strategies[20], however this was not included in our assessment.

### Examination of Identified SVs in CHM1 and NA12878

We examined the full set of identified simple and complex SVs in both CHM1 and NA12878. As expected, we rediscovered many previously reported deletions, duplications and inversions (Table 1). In some cases, we were also able to identify dispersed duplications that were incorrectly identified as overlapping tandem duplication and deletion events in prior reports (Figure 5a, Supplemental Figure 7). Furthermore, we found a recurrence of particular types of CSVs, including inverted-duplication and deletion-inversion events (Figure 5b,c,d, Supplemental Figures 8-10) suggesting that they are likely more common than previously thought. However, there were numerous other CSVs that could not be coalesced into a single classification and may provide future insight into new mechanisms for SV formation.

**Figure 5.**
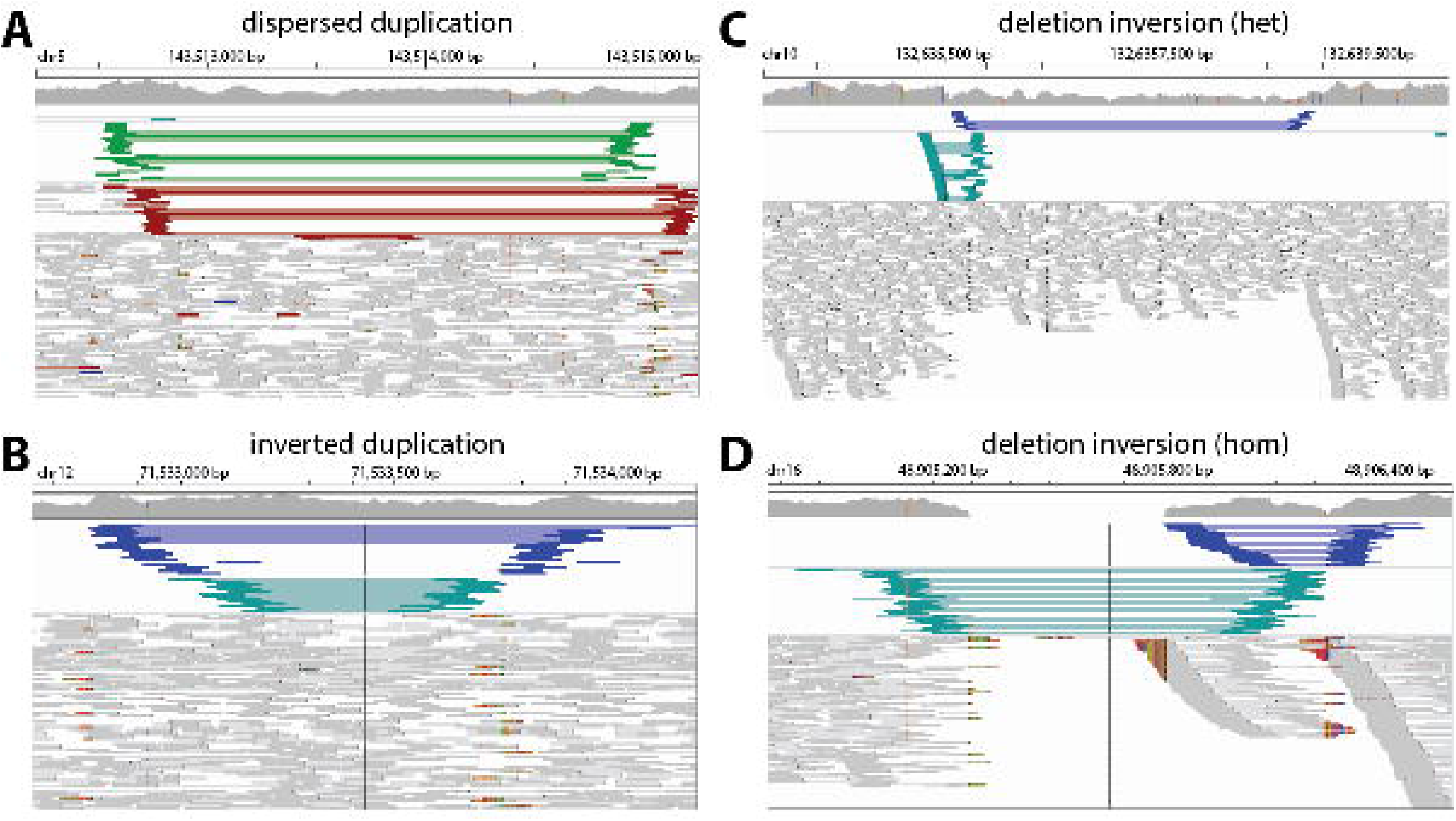
Examples of various types of complex structural variation in NA12878 identified by SVelter. **(A)** IGV screenshot of disperse duplication event predicted by SVelter. Line colors as described in Figure 4. Such regions are typically identified as an overlapping tandem duplication and deletion. **(B)** Example of inverted duplication event. Blue lines in IGV indicated reverse-reverse read pair orientation while dark green lines indicate forward-forward read pair orientation. **(C)** Region with heterozygous inversion and deletion rearrangement. **(D)** Region with homozygous inversion and deletion rearrangement. All regions shown had PacBio sequences consistent with predicted SVelter structures and were misclassified by other approaches (Supplemental Figures 710)

## DISCUSSION

We have described an integrative approach, SVelter, that can identify both simple and complex structural variants through an iterative randomization process. We show that it has an improved or comparable accuracy to other algorithms when detecting deletions, duplications and inversions but has the additional capability of correctly interpreting and resolving more complex genomic rearrangements with three or more breakpoints. Furthermore, SVelter can resolve structural changes on parental haplotypes individually, allowing for the correct stratification of multiple overlapping SVs. SVelter achieves this by forgoing the assumption of specific patterns of read alignment aberrations as associated with individual rearrangements and instead allowing the underlying sequence itself to dictate the most probable structure.

The ability to accurately identify CSVs in whole genome sequence data is a significant advancement, as currently many such regions are either missed or identified as individual errant events. For example, in our investigation of NA12878 we identified many disperse duplications that were previously reported as overlapping deletion and tandem duplication events as well as other simple deletions and inversions that were in fact part of a larger complex rearrangement (Figure 5). Such regions could be, in part, responsible for the observed discrepancies when comparing different SV algorithms with each other as well as other platforms such as array-CGH[21]. Our observations are also consistent with recent findings by the 1000 Genomes Project[19], however their analysis required the use of multiple long-read sequencing technologies including PacBio and Moleculo to interpret the regions while SVelter is able to correctly resolve the regions from short-insert Illumina sequences alone. Although long-read technologies are very well suited for such an application, their use is currently limited to small-scale projects and there have been estimates that over 300,000 genomes will be sequenced using Illumina short-insert reads in 2015 alone. Thus, approaches like SVelter that perform accurately on such data sets are likely to have a larger impact on correctly reporting complex structural genomic aberrations.

One limitation of SVelter is that even with our efficiency enhancements it still exhibits a longer processing time with respect to the other SV algorithms compared here. This is in part due to the randomization strategy but is also owing to the inclusion of a read coverage component, which is not modeled in the other approaches we compared against but contributes to the overall increased accuracy of SVelter. Recent advances have made it possible to analyze a high coverage human genome from sequence to variant calling and annotation in half a day[20], and such applications are very useful for diagnostic applications where speed is a critical component. Nevertheless, the enhanced ability of SVelter to correctly resolve overlapping and complex rearrangements relative to other approaches makes it very useful for projects where the accurate detection of such regions is important. Another limitation of SVelter is that in its current form it has a reduced ability to delineate heterogeneous data, such as commonly found when sequencing cancer genomes. This is due to its expectation of a specific ploidy when iterating between multiple haplotypes. Future work in this area will focus on creating a dynamic structure that can allow different levels of heterogeneity or mosiacism.

## CONCLUSIONS

We have developed and applied a new approach to accurately detect and correctly interpret both simple and complex structural genomic rearrangements. Our comparisons to existing algorithms and data sets show that SVelter is very well suited to identifying all forms of balanced and unbalanced structural variation in whole genome sequencing data sets.

## METHODS

### SVelter Algorithm

SVelter takes aligned Illumina paired-end sequence data in sorted BAM format as input as well as the reference genome against which the sequences were aligned and reports all predicted SVs in both a custom format as well as VCFv4.1. Default parameters are chosen to best balance sensitivity and efficiency, though are adjustable for users to best fit their own data. The SVelter framework consists of three major modules: null model determination, breakpoint detection, random iterative rearrangement, and structure scoring. (Figure 1)

#### Null Model Determination

SVelter first filters the reference genome to exclude regions of low mappability from downstream analysis to increase efficiency by avoiding regions where alignments are unreliable. Such regions include gaps and unknown regions in the reference genome (Ns) and these are integrated with previously reported genomic regions (wgEncodeMapability, obtained from UCSC Genome Browser) that are of low mappability to form a final version of excluded regions. Next, default distributions of insert size (IS), read depth (RD) and physical coverage (PC) are determined by sampling either randomly, or from a set of copy neutral (CN2) genomic regions defined as places in the genome where no polymorphic CNVs, segmental duplications, or repetitive elements have been annotated and thus providing a good estimate of the baseline alignment characteristics[15]. For efficiency, SVelter models these by default as normal distributions with an observed minimal reduction of accuracy on well-behaved sequence libraries, but also provides options for more complex models (i.e. binomial distribution for IS and negative binomial distribution for RD and PC) when the input data is of lower quality.

#### Detection and Clustering of Putative Breakpoints

SVelter next scans the input alignment file to define putative breakpoints (BPs) where the sample genome differs from the reference. These are defined through the identification of aberrant read alignments. Clusters of read pairs (RP) showing abnormal insert length or aberrant mapping orientation may indicate breakpoints nearby, while reads with truncated (clipped) split read (SR) alignments are indicative of precise breakpoint positions. SVelter specifically defines aberrant reads as follows:

1. RPs outside expected insert size (mean ± 3*std)
2. RPs with aberrant pair orientation (FF,RR,RF)
3. SRs with high average base quality (>20) clipped portion with minimum size fraction of overall read length (>10%)

It should be noted that the specific parameters listed were set as default empirically and can be adjusted by the user. Discordant RPs of the within a window of mean insert size + 2*std distance and of the same orientation are clustered together. Next, split reads within this window and downstream of the read direction are collated and the clipped position is considered as a putative breakpoint. If no such reads exist, the rightmost site of forward read clusters or leftmost site of reverse read clusters is assigned instead. For each cluster of aberrant RPs, a BP is assigned if the total number of split reads exceeds 20% of the read depth or the total number of all aberrant reads exceeds 30%. For samples of poorer quality, higher cutoffs might be preferred. Each putative BP will be paired with other BPs that’s defined by mates of its supporting reads. BP pairs that intersect or are physically close (<1kb) to each other will be further grouped and reported as a BP cluster for the next step.

#### Random Iterative Rearrangement

For each BP cluster containing n putative BPs, a randomized iterative procedure is then applied on the n-1 genomic blocks between adjacent BPs. SVelter has three different modules implemented for this step: diploid module (default) that detects structural variants on both alleles simultaneous, heterozygous module that only report high quality heterozygous SVs and homozygous module for high quality homozygous SVs only. For the diploid module, a simple rearrangement (deletion, inversion or insertion) is randomly proposed and applied to each block on one allele while the other allele is kept unchanged and the newly formed structure is scored against the null models of expectation for each feature through the scoring scheme described below. A new structure is then selected probabilistically from the distribution of scores such that higher scores are more likely but not assured. The same approach is then applied to the other allelic structure representing a single iteration overall. For heterozygous and homozygous modules, only one allele is iteratively rearranged while the other allele remains either unchanged or is mirrored, respectively.

The iterative process will terminate and report a final rearranged structure if one of the following situations is met:

1. No changes to a structure after 100 continuous iterations
2. The maximum number of iterations is reached (100,000 as default)

After the initial termination, the structure is reset and the process is repeated for another 100 iterations while avoiding the fixed structure, at which point the highest scoring structure overall is chosen.

#### Structure Scoring

For a rearranged structure, all read pairs originally aligned within the region are scored independently and averaged for the final score. The read pair score (SRP) consists of four parts: Insert Size (SIS), Pair Orientation (SPO), Read Depth (SRD) and physical coverage through a BP (STB) in the following equation:

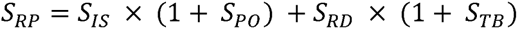

Using the distribution of insert sizes described above, the SIS is calculated as the log transformation of the probability that a particular insert size is observed. RDS is defined in the same way using the observed read depth. SPO is calculated as the faction of read present in the region with aberrant orientation over the total:

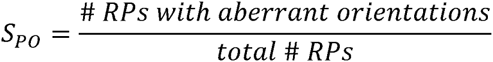

STB is determined using a Chi-square using the physical coverage through a particular breakpoint:

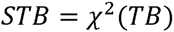

### Performance Assessment

Both simulated and real data were used to evaluate performance of SVelter. To produce simulation datasets, we altered the human GRCh37 reference genome to include both homozygous and heterozygous simple SVs and complex SVs independently while adding micro-insertions and short tandem repeats around the junctions in frequencies consistent with previously reported breakpoint characteristics[22]. Details about specific types of SVs simulated are summarized in Supplementary Table 1. Paired-end reads of 101bp with an insert size of 500bp mean and 50bp standard deviation were simulated using wgsim (https://github.com/lh3/wgsim) across different read depths (10X, 20X, 30X, 40X, 50X).

For comparisons using real sequence data, we adopted two previously published samples: a haploid hydatidiform mole (CHM1)[16] and a well-characterized HapMap/1000 Genomes Project sample (NA12878)[17]. CHM1 has been deep sequenced by Illumina whole-genome sequence to 40X and by Single Molecule, Real-Time (SMRT) sequencing to 54X, and SVs of the sample have been detected and published by the same group as well (http://eichlerlab.gs.washington.edu/publications/chm1-structural-variation/). NA12878, together with the other 16 members from CEPH pedigree 1463, has been deep sequenced to 50X by Illumina HiSeq2000 system (http://www.illumina.com/platinumgenomes/). Additionally, the Genome in a Bottle (GIAB) Consortium has published the PacBio sequencing data (20X) of NA12878 and also provided a set of high-confident SV calls[18].

Three other widely used algorithms: Pindel, Delly and Lumpy were selected for the comparison. We applied SVelter, Pindel and Lumpy to both simulated and real data with default settings, except for that SVelter’s homozygous module was used for CHM1. Delly was run with the insert size cutoff (-s) set as 10 as recommended by the author. All algorithms were run using the same set of genomic regions marked for exclusion and were filtered to only consider SVs >100bp.

#### Assessment of Simulated Simple SVs

For simulated datasets, we compared the performance of each algorithm by calculating their sensitivity and specificity on each type of simple SV (deletion, disperse duplication, tandem duplication, inversion). As Lumpy reports breakpoints in terms of range, we calculated the median coordinate of each reported interval and consider it as the breakpoint for downstream comparison. A reported SV would be considered as a true positive (TP) if the genomic region it spanned overlapped with a simulated SV of the same type by over 50% reciprocally. As Delly and Lumpy didn’t differentiate tandem and dispersed duplication in their SV report, we compare their reported duplications to both simulated tandem and dispersed duplications independently to calculate sensitivity, but use the entire set of simulated duplications together for the calculation of specificity. In this manner, the result will be biased towards higher TP and TN rates for these approaches. Dispersed duplications reported by Pindel were very rare and as such were processed in the same way as Delly and Lumpy.

#### Assessment of Real SVs

We initially made use of reported simple and complex SVs in CHM1 and NA12878 as gold standard sets, however the FP rate of each algorithm were high compared to previously published performance. To augment this set, we therefore have developed our own approach to validate simple and complex SVs using PacBio long reads. For each reported SV, we collect all PacBio reads that go through the targeted region and hard clip each read prior to the start of the region. We then compare each read to the local reference and an altered reference reflecting the structure of the reported SV by sliding a 10bp window through the PacBio read and aligning it against the reference sequence. Coordinates of each window are plotted against its aligned position in the form of a dotplot. Theoretically speaking, if a read was sampled from the reference genome, a diagonal line should be observed. However, if a read was sampled from an altered genomic region, a continuous diagonal line would only show when plotted against a correctly resolved sequence. In this manner, shorter SVs (<5kb) can be validated by accessing the deviation of all dots from diagonal. For each PacBio Read, the score:

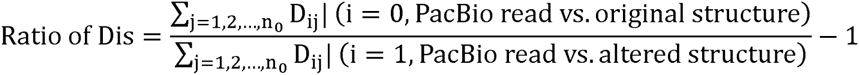

is assigned, so that a positive Ratio of Dis indicates the priority of altered genome over reference genome, and vise versa. The validation score of an SV is integrated from all PacBio reads spanning through it using an indicator function:

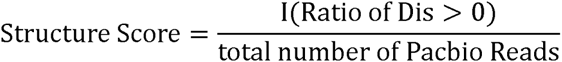

SVs with validation score >0.5 for haploid genome, or >0.3 for diploid genome would be considered validated.

For longer (>5kb) SVs, PacBio reads spanning through the whole targeted region are rarely observed. In this situation, we scored each breakpoint by adding 500bp flanks and assessing each individually. The final validation score is then determined through the collation of matches from all breakpoints.

We reassessed our initial true positive (TP) and false positive (FP) simple calls from each algorithm by combining our PacBio validated SVs from each algorithm together with the reported calls. For simple SVs, we utilized a 50% reciprocal overlap criterion. However, for CSVs we utilized a more complex comparison strategy to take into account that some algorithms will often detect individual parts of a complex rearrangement as distinct events. With each CSV predicted by SVelter, we extracted SVs with over 50% reciprocal overlap from other algorithms and calculated the validation score for each of them using our PacBio validation approach described above. When multiple SVs were extracted from an algorithm, averaged scores were adopted. Validation scores of a CSV from all algorithms were ranked and normalized from 0 to 1 for comparison.

## SOFTWARE AND DATA AVAILABILTY

The software package *SVelter* is available for download at https://github.com/millslab/svelter, and additional documentation regarding specific software usage and parameters, supporting files, algorithm comparisons and simulated data sets are provided at this site.

Sequence data used in this analysis were obtained from the following resources: CHM1 – Resolving the complexity of the human genome using single-molecule sequencing (http://eichlerlab.gs.washington.edu/publications/chm1-structural-variation/) [16]

NA12878 – Genome in a Bottle Consortium (https://sites.stanford.edu/abms/giab)[17], Illumina Platinum Genomes (http://www.illumina.com/platinumgenomes/)

## AUTHOR CONTRIBUTIONS

XZ developed and implemented the algorithms and wrote the source code. REM conceived the analytical framework, devised the experiments and supervised the project. SBE, BM, and JMK performed the PCR validation experiments. REM and XZ prepared the figures and wrote the manuscript. All authors read and approved of the final manuscript.

## DESCRIPTION OF ADDITIONAL DATA FILES

The following additional data are available with the online version of this paper. Additional data file 1 contains Supplemental Table 1 outlining the type and number of SVs included in each simulated genome and the stratified results for each algorithm. Additional data file 2 contains Supplemental Tables 1-3 and Supplemental Figures 1-10. Additional data file 3 additional Supplemental Methods outlining the software and parameter usage that was used to generate the presented results.

## ACKNOWLEDGMENTS

This project was supported in part through funds from the University of Michigan, the NIH/NHGRI (1R01-HG007068-01A1), and NIH/Common Fund (DP5OD009154).

